# Inactivation of the *Fusobacterium nucleatum* Rnf complex reduces FadA-mediated amyloid formation and tumor development

**DOI:** 10.1101/2025.04.03.647037

**Authors:** Timmie A. Britton, Ju Huck Lee, Chungyu Chang, Aadil H. Bhat, Yi-Wei Chen, Rusul Mohammed Ali, Chenggang Wu, Asis Das, Hung Ton-That

## Abstract

The Gram-negative anaerobe *Fusobacterium nucleatum* is an oral oncobacterium that promotes colorectal cancer (CRC) development with the amyloid-forming cell surface adhesin FadA integral to CRC tumorigenesis. We describe here molecular genetic studies uncovering a novel mode of metabolic regulation of FadA-mediated tumor formation by a highly conserved respiratory enzyme known as the Rnf complex. First, we show that genetic disruption of Rnf, via *rnfC* deletion, significantly reduces the level of *fadA* transcript, accompanied by a near-complete abolishment of the precursor form of FadA (pFadA), reduced assembly of FadA at the mature cell pole, and severe defects in the osmotic stress-induced formation of FadA amyloids. We show further that the Rnf complex regulates three response regulators (CarR, ArlR, and S1), which modulate the expression of pFadA, without affecting *fadA* transcript. Consistent with our hypothesis that these response regulators control factors that process FadA, deletion of *rnfC*, *carR*, *arlR*, or *s1* each impairs expression of the signal peptidase gene *lepB*, and FadA production is nearly abolished by CRISPR-induced depletion of *lepB*. Importantly, while *rnfC* deletion does not affect the ability of the mutant cells to adhere to CRC cells, *rnfC* deficiency significantly diminishes the fusobacterial invasion of CRC cells and formation of spheroid tumors *in vitro*. Evidently, the Rnf complex modulates the expression of the FadA adhesin and tumorigenesis through a gene regulatory network consisting of multiple response regulators, each controlling a signal peptidase that is critical for the post-translational processing of FadA and surface assembly of FadA amyloids.

**IMPORTANCE:** The *<u>R</u>hodobacter* <u>n</u>itrogen-fixation (Rnf) complex of *Fusobacterium nucleatum* plays an important role in the pathophysiology of this oral pathobiont, since genetic disruption of this conserved respiratory enzyme negatively impacts a wide range of metabolic pathways, as well as bacterial virulence in mice. Nonetheless, how Rnf deficiency weakens the virulence potential of *F. nucleatum* is not well understood. Here, we show that genetic disruption of the Rnf complex reduces surface assembly of adhesin FadA and FadA-mediated amyloid formation, via regulation of signal peptidase LepB by multiple response regulators. As FadA is critical in the carcinogenesis of colorectal cancer (CRC), the ability to invade CRC cells and promote spheroid tumor growth is strongly diminished in an Rnf-deficient mutant. Thus, this work uncovers a molecular linkage between the Rnf complex and LepB-regulated processing of FadA – likely via metabolic signaling – that maintains the virulence potential of this oncobacterium in various cellular niches.

## INTRODUCTION

The *<u>R</u>hodobacter* <u>n</u>itrogen fixation (Rnf) complex is a membrane bound respiratory enzyme that catalyzes the oxidation of reduced ferredoxin and the reduction of NAD+, thereby establishing an ion-motive force and permitting substrate import and ATP biosynthesis (1–4). Originally discovered in the phototrophic bacterium *Rhodobacter capsulatus* for its role in nitrogen fixation (1), the Rnf complex is highly conserved in Gram-positive and Gram-negative bacteria including significant pathogens, with multiple genes coding for the Rnf subunits clustered together into an operon (4); for example, the *rnfABCDGEH* cluster of *R. capsulatus* (5), the *rnfABCDGE* of *Escherichia coli* (6), and the *rnfCDGEAB* cluster of *Clostridium tetani* and *Fusobacterium nucleatum* (7, 8). Although the role of the Rnf complex in energy conservation has been well documented (4, 9–11), its role in bacterial pathogenesis has begun to emerge only recently. In the Gram-positive gut bacterium *Clostridium sporogenes*, a *rnfB* mutant is shown to be attenuated for growth in the mouse gut (12), while in the Gram-negative oral anaerobe *F*. *nucleatum*, a mutant devoid of *rnfC* (Δ*rnfC*) displays virulence defects in a mouse model of preterm birth (8). In Δ*rnfC* cells, catabolism of many amino acids, including cysteine, histidine and lysine, is significantly affected, hence reducing production of ATP and many metabolites such as hydrogen sulfide and butyrate (8). Note that although the fusobacterial *rnfC* mutant exhibits slow growth and cell morphological defects, the mutant’s colony forming ability (CFUs) is comparable to that of the parent strain (8). Thus, the attenuated virulence of the Rnf mutant may be independent of a reduction in bacterial burden during infection. While the role of the *C*. *sporogenes* Rnf complex in metabolism has been implicated in gut colonization (12), how genetic disruption of the Rnf complex affects *F. nucleatum* virulence remains to be elucidated.

*F*. *nucleatum*, an opportunistic pathogen commonly present in the oral cavity of healthy individuals, is associated with several extra-oral pathologies, including adverse pregnancy outcomes such as preterm birth and neonatal sepsis (13, 14) and the promotion of colorectal cancer (CRC) (15–18). Among several adhesins that are implicated in the interaction of *Fusobacterium* with its host, FadA plays a crucial role in fusobacterial pathogenesis. A mutant lacking *fadA* is defective in placental colonization and stimulation of CRC cell growth (18, 19). Through FadA and E-cadherin, *F. nucleatum* induces expression of Annexin A1 in cancerous cells, with Annexin A1 acting as a modulator of Wnt/ β-catenin signaling (20). Curiously, the FadA protein exists in two forms – the full-length precursor form (pFadA) of 129 residues and the secreted mature FadA (mFadA) missing its signal peptide but migrating slower than pFadA on SDS-PAGE (21, 22). An active FadA complex (FadAc), comprised of both mFadA and pFadA, is necessary and sufficient to promote CRC cell growth via FadA binding to E-cadherin on CRC cells (18). Importantly, under stress and disease conditions, FadA also forms amyloid-like structures on the bacterial surface – involving pFadA-assisted crosslinking of FadA filaments – that are critical for CRC progression in mice (23). It is noteworthy that the secretion of FadA requires a Fap2-like autotransporter (23), with Fap2 previously shown to mediate fusobacterial binding to Gal-GalNAc, a biomarker that is abundantly expressed by adenocarcinomas (24).

RadD is another adhesin that has recently been shown to promote fusobacterial binding to CRC cells (25). Originally identified as the major coaggregation factor that mediates fusobacterial adherence to many oral bacteria (26, 27), RadD directly binds to CD147, a glycoprotein that is highly expressed on the surface of tumor cells, triggering a PI3K-AKT-NF-κB-MMP9 signaling reaction to enhance tumorigenesis (25). These findings suggest that *F. nucleatum* may utilize multiple pathways to promote tumorigenesis and CRC progression. It is noteworthy that the RadD-mediated coaggregation of fusobacteria with other oral bacteria is diminished when the *F. nucleatum* Rnf complex is disrupted via deletion of *rnfC* (27). This defect is not caused by a reduced bacterial surface expression of RadD, but rather due to global metabolic defects leading to an excessive accumulation of environmental lysine, which binds to RadD and in turn, inhibits RadD-mediated bacterial binding to oral bacteria (27). Yet another notable effect of disrupting the Rnf complex is the diminished expression of *megL*, which encodes the L-methionine γ-lyase MegL involved in methionine/cysteine metabolism (28). To date, how the genetic disruption of the Rnf complex affects global gene expression in *F. nucleatum* has not been elucidated.

Here, we show that genetic disruption of the Rnf complex, via *rnfC* deletion, decreases surface expression of FadA and FadA-mediated amyloid formation under osmotic stress, via an indirect regulation by several response regulators. In fact, these regulators control the expression of the signal peptidase LepB that processes the precursor form of FadA. As FadA plays an important role in the carcinogenesis of CRC, inactivation of Rnf reduces fusobacterial ability to invade CRC cells and promote tumor growth. Our study presented here reveals a molecular linkage between the Rnf complex and gene regulation – likely via metabolic signaling – that modulates *F. nucleatum* virulence.

## RESULTS

### Genetic disruption of the Rnf complex reduces FadA expression and FadA amyloidogenesis on the fusobacterial cell surface

In our previous work, we have uncovered a major role of the Rnf complex in fusobacterial metabolism, gene expression, and pathogenesis by demonstrating that the genetic disruption of the complex causes significant defects in each of these processes (8). To better characterize the impact of Rnf-dependent metabolism on fusobacterial virulence, we examined whether disrupting the Rnf complex with a deletion of *rnfC* affects the expression of FadA and Fap2, two fusobacterial adhesins that are important for fusobacterial colonization and CRC promotion (14, 20, 24). To analyze FadA, protein samples from whole-cell lysates of parent, mutant, and complemented *F. nucleatum* strains were examined by western blotting using a polyclonal antibody against FadA (α-FadA) that detects both precursor (pFadA) and mature (mFadA) forms (22). Remarkably, as compared to both the parent and complemented strains, deletion of *rnfC* (Δ*rnfC*) drastically reduced the level of pFadA, coupled with an increased level of mFadA, which migrated slower than pFadA according to the previous assignments of these protein forms (22, 23) (Fig. 1A). To examine whether the reduced level of pFadA was due to a reduction in the expression of *fadA* transcripts, we extracted RNA from these samples and performed quantitative reverse transcription polymerase chain reaction (qRT-PCR) analysis. Consistently, there was a 10-fold reduction of *fadA* mRNA level in the Δ*rnfC* mutant, relative to the parent strain (Fig. 1B). Since FadA is a cell surface protein (23), we next probed for the cell surface level of FadA by immunofluorescence microscopy (IFM), whereby fusobacterial cells were first stained with α-FadA antibody and then with Alexa488-conjugated IgG, together with DAPI staining the chromosome. Consistent with the above results, the FadA signal of the Δ*rnfC* mutant was significantly reduced compared to that of the parent strain, and this defect was rescued by ectopic expression of FadA (Fig. 1C & 1D). Note that the surface localization of FadA was restricted to the cell pole (Fig. 1C).

**Figure 1:**
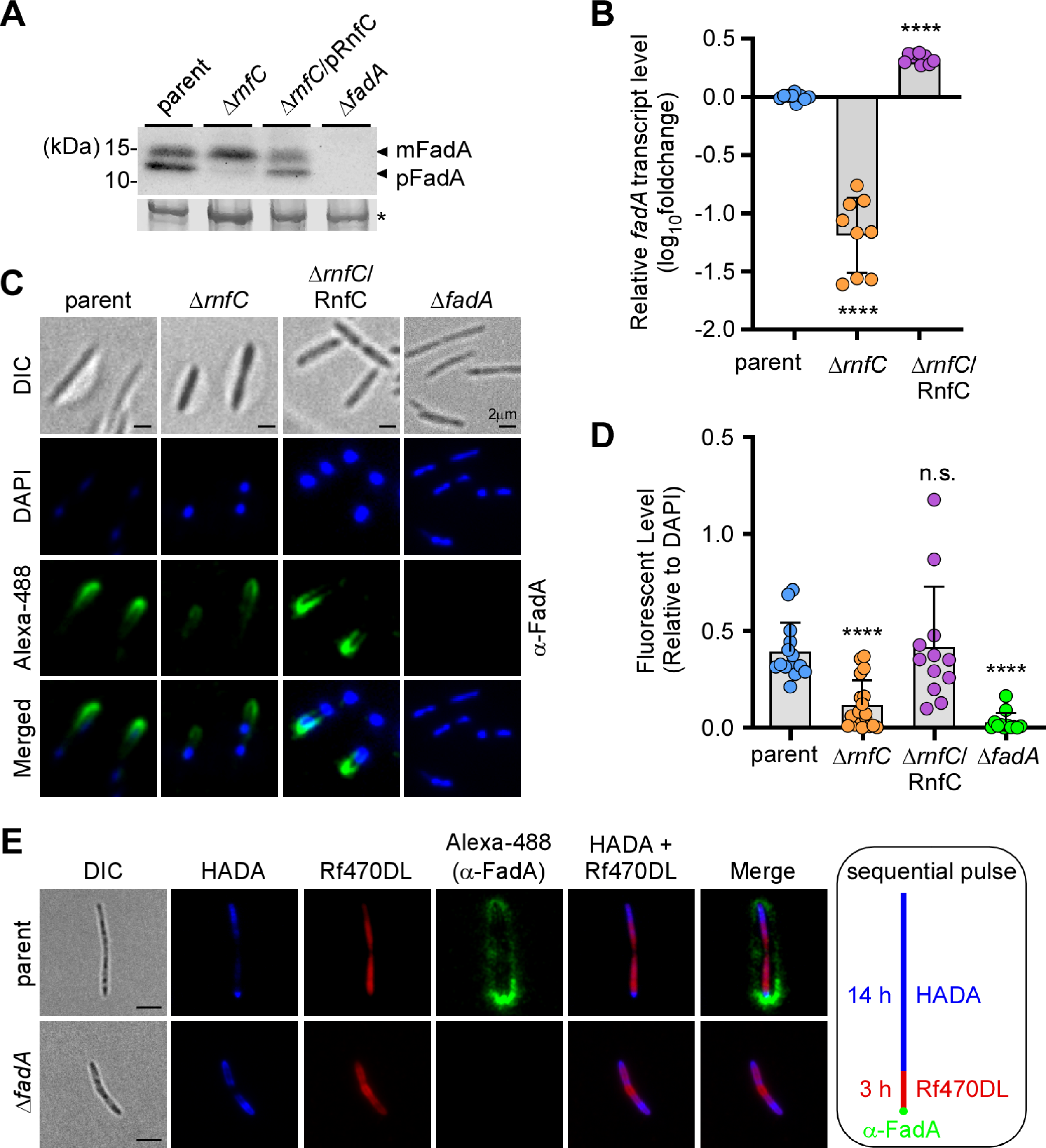
Genetic disruption of the Rnf complex, via *rnfC* deletion, reduces expression of FadA. **(A)** Protein samples obtained from whole-cell lysates of normalized cultures from indicated fusobacterial strains were subjected to immunoblotting with antibodies against FadA (α-FadA). Black arrows mark the precursor form of FadA (pFadA) and mature FadA (mFadA), with a Coomassie Blue stained band (*) from the same blotting membranes used as a loading control. **(B)** Normalized overnight cultures of indicated strains were used to isolate total RNA for qRT-PCR to determine the transcript levels of *fadA*. All qRT-PCR data was normalized by 16s rRNA transcript abundance for each sample. **(C)** Overnight cultures of indicated strains were first stained with α-FadA, followed by Alexa488-conjugated secondary antibodies (green), as well as DAPI (blue). Surface localization of FadA was visualized by a fluorescence microscope, and representative images are shown. **(D)** FadA signal relative to DAPI signal from indicated strains shown in panel C was quantified and presented. **(E)** Cells of parent and Δ*fadA* mutant strains grown to log-phase were harvested and sequentially labeled with fluorescent dyes HADA and Rf470DL for 14 h and 3 h, respectively, followed by labeling with α-FadA and Alexa488 as described in panel C. Samples were analyzed by fluorescence microscopy. All results were obtained from three independent experiments performed in triplicate. Significance was calculated by a Student’s t-test; **** P < 0.0001.

To uncover the spatiotemporal dynamics of FadA deposition on the fusobacterial cell surface, we employed fluorescently-tagged D-amino acids (FDAAs), which are covalently incorporated into the bacterial cell wall by endogenous transpeptidases without disrupting bacterial metabolism or cell growth; as such, these compounds are used to probe peptidoglycan biosynthesis (29, 30). In this experiment, fusobacterial cells were first cultured in the presence of HCC-amino-D-alanine (HADA) for 14 h prior to pulse-labeling with Rf470DL for 3 h.

Fusobacteria were then fixed for immunostaining with α-FadA and Alexa488-conjugated IgG (Fig. 1E). As shown in Fig. 1E, HADA labeled oldest peptidoglycan at the cell poles of two dividing cells, while Rf470DL marked newly synthesized peptidoglycan. Curiously, strong FadA signal was mainly observed at the old cell pole, although the signal was also observed along the cell envelope, albeit with much lower intensity (Fig. 1E).

Next, using the same set of samples described above, we analyzed surface expression of the outer membrane protein Fap2 by immunoblotting with antibodies against Fap2 (α-Fap2). To our surprise, the expression of Fap2 was significantly increased in the Δ*rnfC* mutant, relative to the parent and complement strains (Fig. 2A & 2B). Consistent with the western blot analysis, deletion of *rnfC* also resulted in increased cell surface signal for Fap2 that appeared to be punctate (Fig. 2C & 2D).

**Figure 2:**
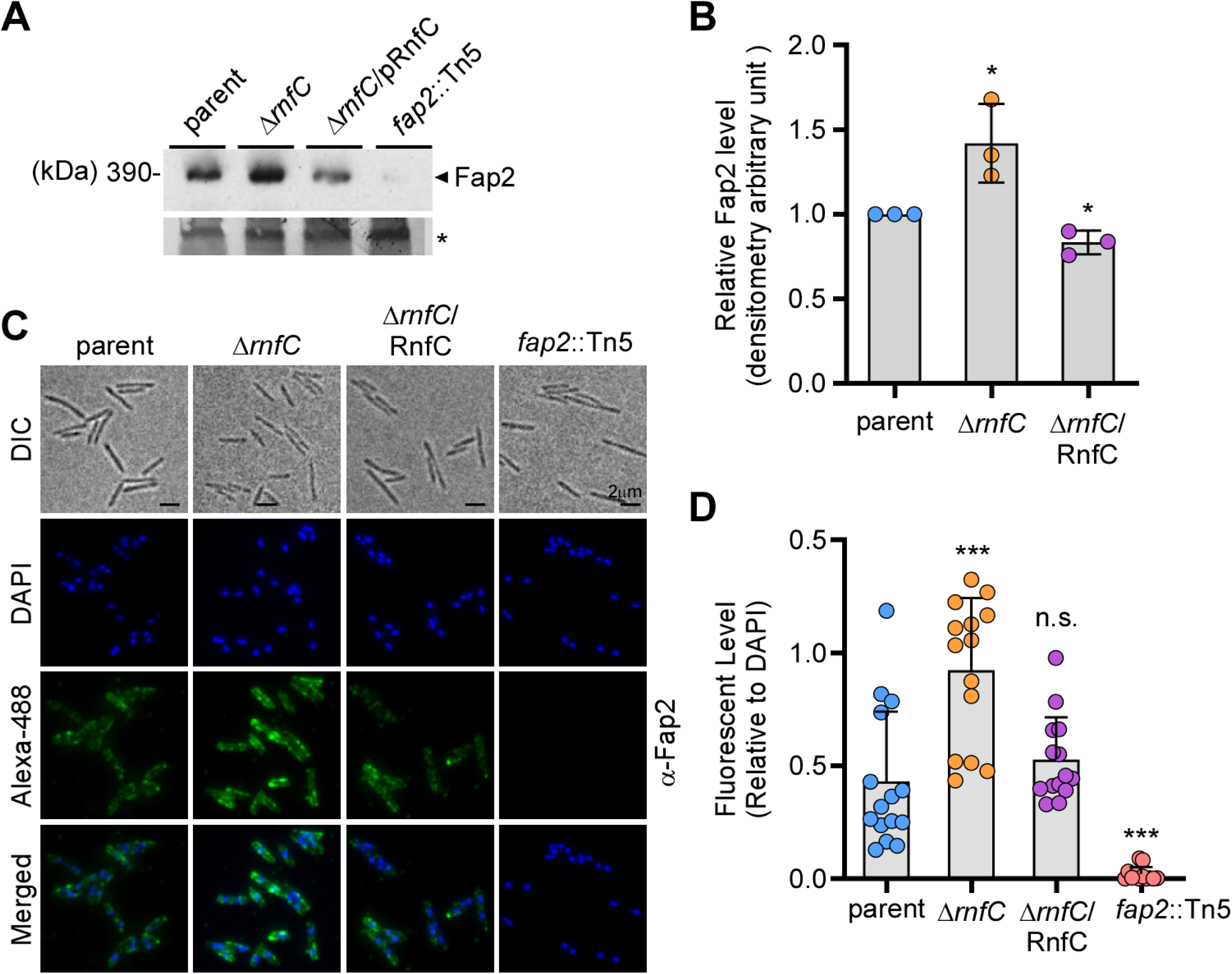
Deletion of *rnfC* increases expression of Fap2. **(A)** Protein samples obtained from whole-cell lysates of normalized cultures from indicated strains grown overnight were subjected to immunoblotting with antibodies against Fap2 (α-Fap2), with a Coomassie Blue stained band (*) used as a loading control. **(B)** Fap2 signal of indicated strains from three independent experiments in panel A was quantified by densitometry, with the Coomassie Blue stained band used as control. **(C)** Overnight cultures of indicated strains were stained with α-Fap2, followed by Alexa488-conjugated secondary antibodies (green), as well as DAPI (blue). Surface localization of Fap2 was visualized by a fluorescence microscope. **(D)** Fap2 signal, relative to DAPI signal from indicated strains in panel C, was quantified by ImageJ. All results were obtained from three independent experiments performed in triplicate. Significance was calculated by a Student’s *t*-test; * *P* ≤ 0.05; ** *P* < 0.01.

Given that under stress conditions FadA forms amyloids on the cell surface that can be visualized by IFM using antibodies raised against human amyloid beta 42 (α-Aβ42) (23), we performed a similar experiment and found that *F. nucleatum* grown in media with high salt (100 mM NaCl) produced markedly more FadA amyloids than cells grown without added NaCl (Fig. 3A). Consistent with our western blot and immunofluorescent results for FadA described above (Fig. 1), the cell surface associated FadA-amyloids we detected with α-Aβ42 were significantly reduced in the Δ*rnfC* mutant, as compared to the parent and complemented strains (Fig. 3B & 3C). Together, the results indicate that the disruption of the RnfC complex, via *rnfC* gene deletion, causes a significant reduction in *fadA* gene expression, which in turn reduces the level of FadA protein and the formation of FadA amyloids on the cell surface, while also increasing expression of the outer membrane adhesin Fap2.

**Figure 3:**
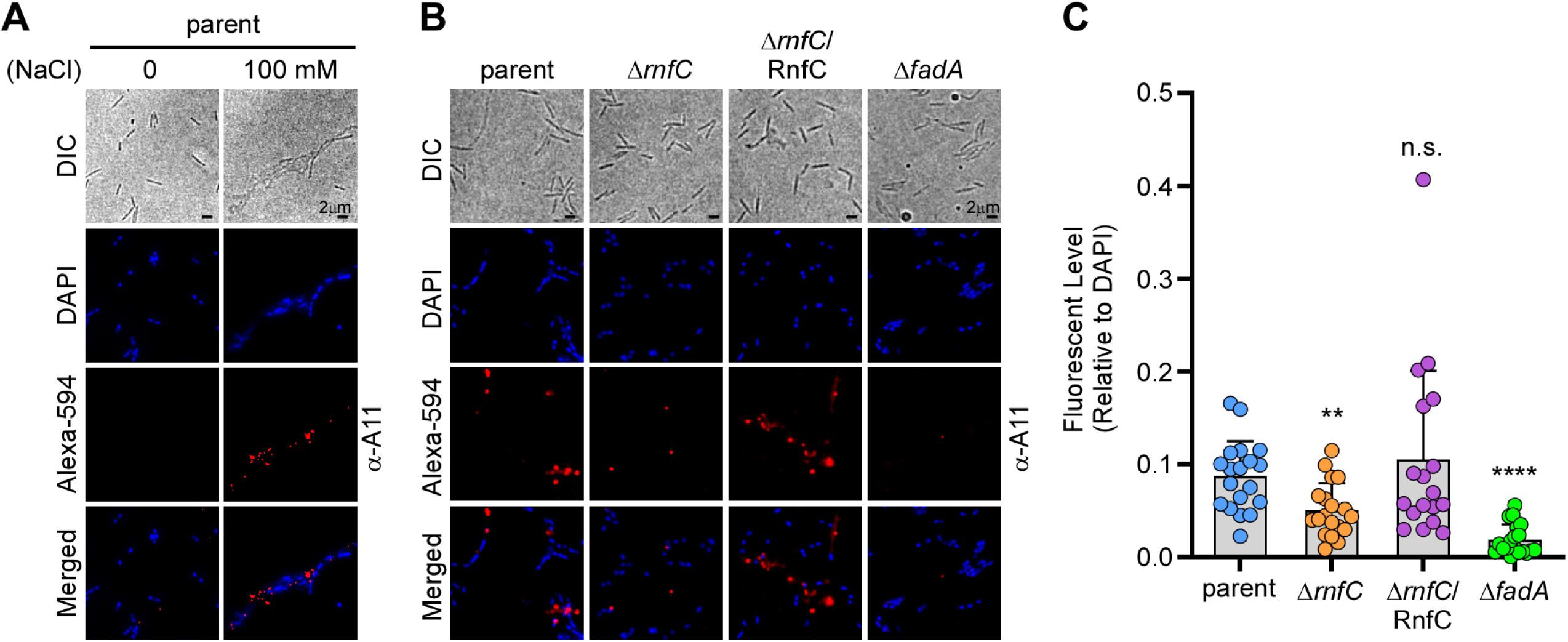
Genetic disruption of the Rnf complex reduces formation of FadA-mediated amyloids. **(A)** Parental cells grown overnight in the presence or absence of 100 mM NaCl were harvested for immunofluorescence microscopy. Cells were first stained with antibodies against human amyloid beta 42 (α-A11), followed by staining with Alexa594-conjugated secondary antibodies (red), as well as DAPI (blue), prior to microscopic analysis. **(B-C)** A similar experimental procedure was performed with the indicated strains. Quantification of amyloid signal, via Alexa594, relative to DAPI signal in these strains is shown in panel C. All results were obtained from three independent experiments performed in triplicate. Significance was calculated by a Student’s *t*-test; **, *P* < 0.01 and ****, *P* < 0.0001.

### FadA expression is modulated post-transcriptionally by several two-component system response regulators

We next investigated how genetic disruption of the Rnf complex affects *fadA* gene expression. Prompted by our previous observation that the response regulator CarR of the two-component system (TCS) CarRS regulates *megL* expression in *F. nucleatum* (27), we proceeded to determine whether *fadA* is similarly regulated. By bioinformatics analysis, we identified 7 TCS’s in *F. nucleatum* including ModRS, CarRS, and ArlRS (see Table S1) (27, 31, 32). The response regulator CarR modulates a large regulon including *megL*, *radD*, and many lysine metabolic genes (27), whereas ModR regulates genes coding for factors involved in oxidative stress, metabolism, and many other processes (31). The targets of ArlR have not been described so far, although its homolog in *Staphylococcus aureus* controls many genes involved in adhesion, autolysis, and proteolysis (33). Other response regulators of *F. nucleatum* are predicted to be homologous to CheY, EutV, S1, and YpdB (Table S1). To examine whether any of these response regulators control FadA expression, we generated corresponding non-polar, in-frame deletion mutants and analyzed FadA protein and mRNA levels in these strains by western blotting and qRT-PCR, respectively. Remarkably, a significant reduction in the level of FadA protein was observed in each of three different response regulator mutants (Fig. 4A-4B), while none of the mutants significantly diminished the level of *fadA* transcripts (Fig. 4C). Our finding of a different level of mRNA and protein expression led us to hypothesize that FadA might be subject to some form of post-translational regulation that involves the response regulators CarR, ArlR, and S1.

**Figure 4:**
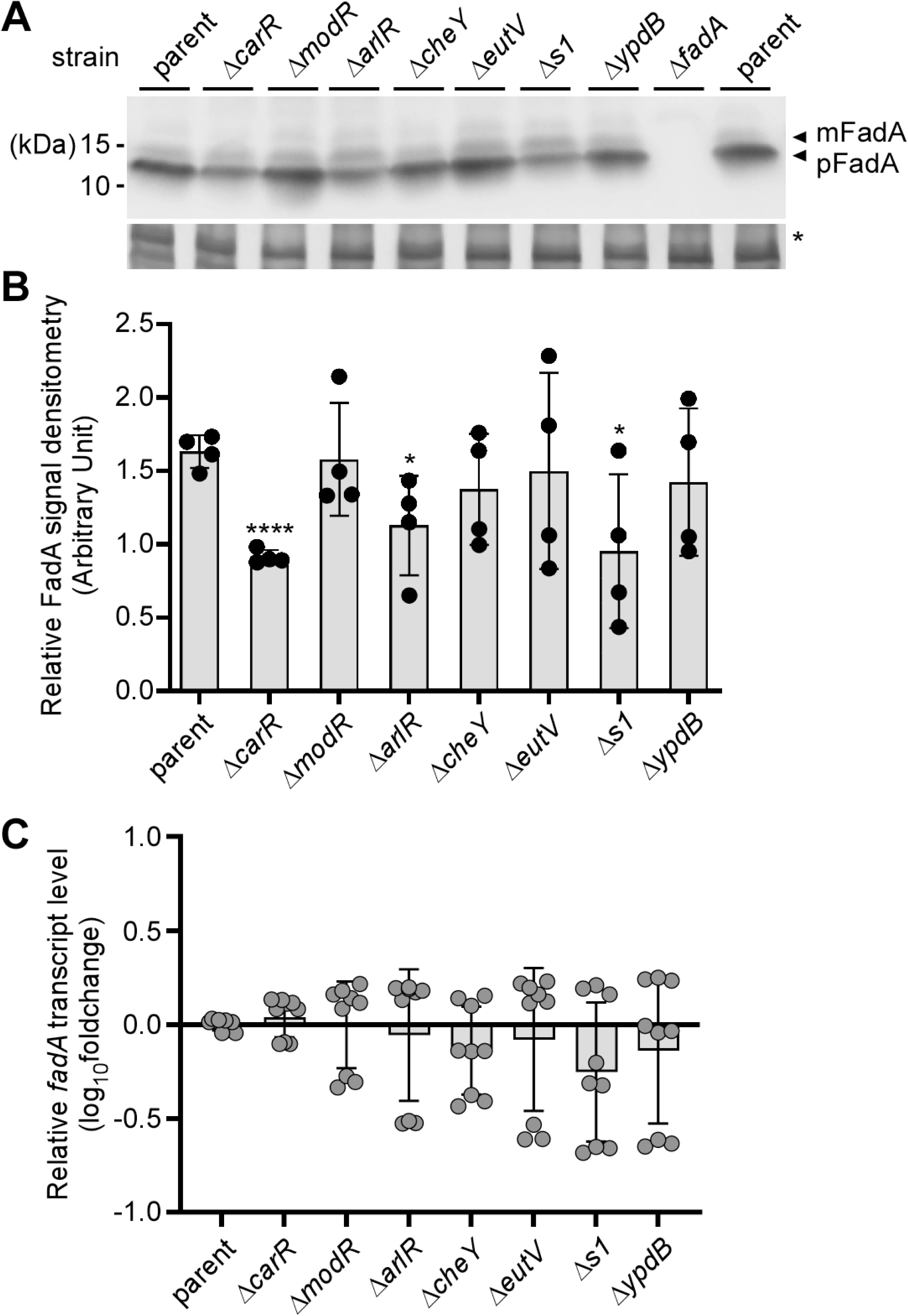
Expression of FadA is modulated by several response regulators of fusobacterial two-component systems. **(A)** Protein samples obtained from whole-cell lysates of normalized overnight cultures of indicated strains were subjected to immunoblotting with antibodies against α-FadA. Arrows mark pFadA and mFadA, with a Coomassie Blue stained band (*) used as a loading control. **(B)** FadA signal of strains in panel A was quantified from three independent experiments by densitometry, with the Coomassie Blue stained band used as control. **(C)** Normalized overnight cultures of indicated strains were used to isolate total RNA for qRT-PCR to determine the expression level of *fadA*, with 16s rRNA transcript abundance used as control. All data was obtained from three independent experiments performed in triplicate. Significance was calculated by a Student’s *t*-test (A) or Mann-Whitney U test (C) according to data distribution; *, *P* ≤ 0.05 and ****, *P* < 0.0001.

### The Rnf complex is required for the expression of the seven response regulator-**encoding genes and signal peptidase *lepB*.**

We previously postulated that a metabolic blockage caused by the deletion of the Rnf complex may trigger gene expression responses from the TCSs of *F. nucleatum* (8). To test this possibility, we measured the transcript levels of these response regulators in the presence or absence of *rnfC* using qRT-PCR. To our astonishment, the mRNA level of all seven response regulators was strongly reduced in the *rnfC* mutant as compared to the parent strain, and complementation of the mutant with a plasmid encoding *rnfC* rescued this defective gene expression (Fig. 5A). Logically, through a loss of the seven response regulators, the depletion of the Rnf complex is expected to have a global deficit in gene expression involving many genes normally targeted by the response regulators. One or more of these undefined genes might be involved in the post-translational regulation of FadA that we postulated above.

**Figure 5:**
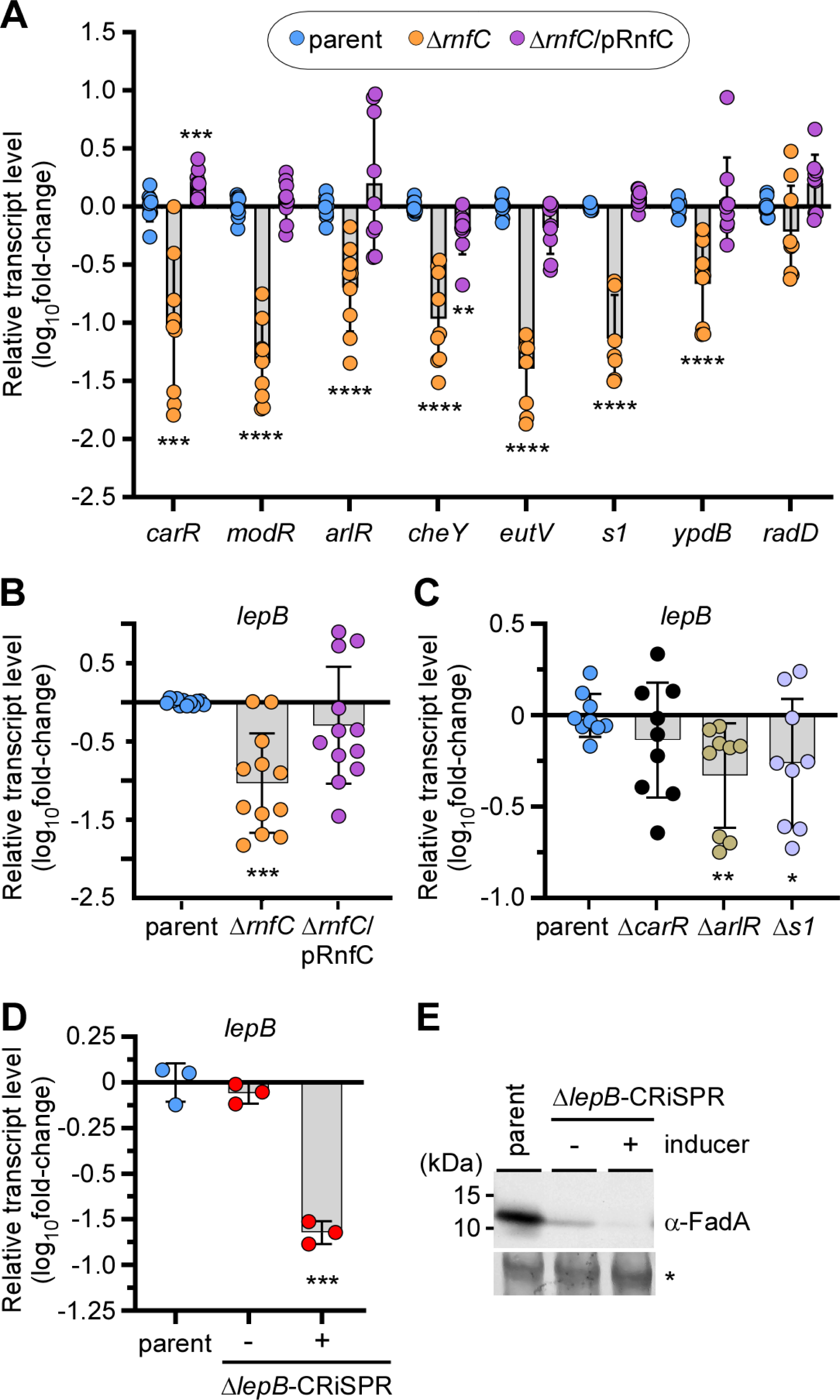
The Rnf complex modulates LepB-regulated cleavage of FadA. **(A-C)** Equivalent overnight cultures of indicated strains were used to isolate total RNA for qRT-PCR to determine the expression levels of *carR, modR, arlR, cheY, eutV, s1, ypdB, and radD* (panel A), as well as *lepB* (panels B-C), with 16s rRNA used as control. **(D-E)** The parent and CRISPRi strains were grown overnight in the presence (+) or absence (-) of 2 mM theophylline (inducer). Normalized cells were harvested for RNA extraction and qRT-PCR, with 16s rRNA used as control (panel E). In a parallel experiment, normalized cells were used for preparation of whole-cell lysates for immunoblotting with α-FadA. A Coomassie Blue stained band (*) was used as a loading control (panel E). All data was obtained from three independent experiments performed in triplicate. Significance was calculated by a Mann-Whitney U test (A) or Student’s *t*-test (B-D) according to data distribution; * *P* < 0.05; ** *P* < 0.01; *** *P* < 0.001; **** *P* < 0.0001.

In search for a post-translational step for regulating the outer membrane protein FadA, we turned our attention to its primary structure for features of inner membrane processing during protein secretion. Indeed, as FadA contains a signal peptide sequence (22) that may be cleaved by the signal peptidase LepB, we next determined whether *rnf* deletion also affects *lepB* expression. Compared to the parent and rescued strains, the *rnfC* mutant expressed a reduced level of *lepB* (Fig. 5B). Because the response regulators CarR, ArlR, and S1 might modulate the expression of FadA without affecting its transcript level (Fig. 4), we further examined by qRT-PCR whether these regulators controls expression of *lepB*. As shown in Fig. 5C, although deletion of *carR* did not significantly reduce the *lepB* transcript level, deletion of *arlR* or *s1* decreased expression of *lepB*.

To determine whether LepB, an essential protein of *F. nucleatum* (34), is the major signal peptidase that processes FadA, we next engineered a conditional knock-out of *lepB* by employing a recently developed gene-editing tool based on CRISPR (35). Strikingly, when the CRISPR system was induced to inactivate *lepB*, we observed a multifold reduction of the *lepB* transcript (Fig. 5D), concomitant with the near abolishment of FadA protein expression (Fig. 5E). Note that FadA was significantly reduced in the non-induced CRISPR condition, most likely caused by leaky expression of CRISPR machinery; as a result of LepB deficiency, the unprocessed precursors of unfolded FadA (pFadA) might be subject to proteolytic quality control (36). Together, these results establish a molecular linkage between the Rnf complex and the gene regulatory cascade that modulates the level of the essential signal peptidase LepB, which in turn processes FadA post-translationally for secretion through the inner and the outer membranes of *F. nucleatum*.

### The Rnf complex promotes fusobacterial invasion of cancer cells and tumor formation *in vitro*

As the FadA adhesin is critical for fusobacterial binding, invasion, and proliferation of CRC cells (18, 20), it was important to determine whether the reduced expression of FadA in the *rnfC* mutant impairs its function as an oncobacterium, i.e. its ability to adhere, invade, and/or induce tumor formation by CRC cells. Therefore, we subjected the aforementioned *F. nucleatum* strains to adherence/invasion and spheroid formation assays as previously reported (31, 37). For measuring bacterial adherence, the CRC HCT116 cells were incubated with fusobacteria at a multiplicity of infection (MOI) of 50 for 4 h and washed prior to CRC cell lysis for enumeration of fusobacterial colony forming units (CFUs), whereas for measuring invasion, the infected HCT116 cells were treated with gentamicin prior to washing and lysing CRC cells for bacterial enumeration. The results showed that while *rnfC* deletion did not affect fusobacterial adherence to CRC cells (Fig. 6A), the mutation significantly reduced fusobacterial invasion of CRC cells, and this defect was rescued by ectopic expression of *rnfC* (Fig. 6B).

**Figure 6:**
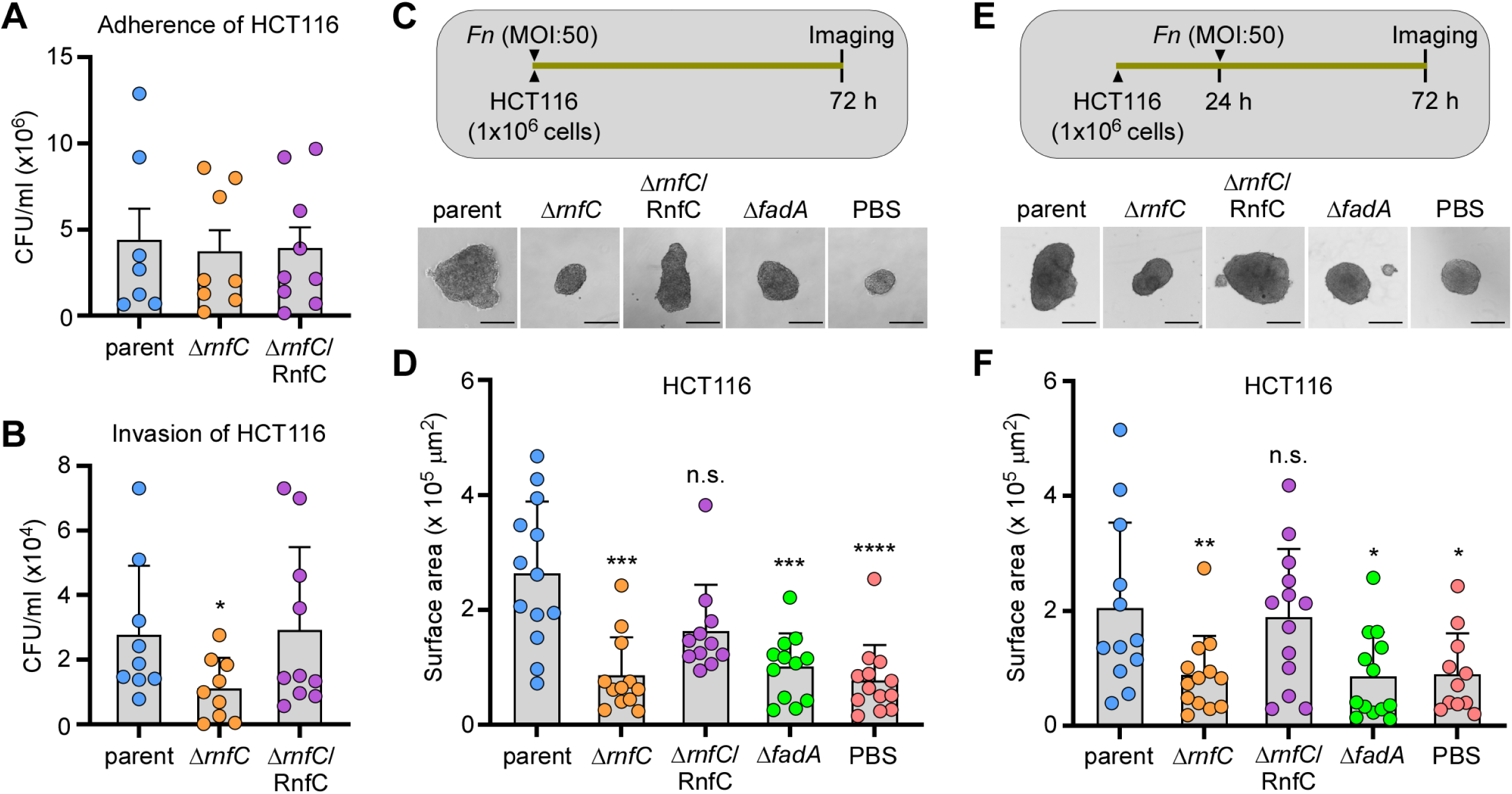
Genetic disruption of the Rnf complex reduces bacterial invasion of cancer cells and tumor formation. **(A)** HCT116 cells were infected with indicated fusobacterial strains at an MOI of 50 for 4 h before being washed off unadhered bacteria and lysed for bacterial enumeration as colony forming units per mL (CFU/mL). **(B)** A similar procedure as A was performed, except that HCT16 cells were treated with 200 µg/mL gentamycin before lysis for bacterial enumeration. **(C-D)** HCT116 cells were infected with indicated strains at an MOI of 50 for 72 h, with PBS used as control, and the resulting spheroids were microscopically analyzed (panel C) and quantified (panel D). **(E-F)** HCT116 cells were allowed to form spheroids for 24 h and treated with indicated strains an MOI of 50. The resulting spheroids were microscopically analyzed (panel E) and quantified (panel F). All results were obtained from three independent experiments performed in triplicate. Significance was calculated by a Mann-Whitney U test; * *P* ≤ 0.05; ** *P* < 0.01; *** *P* < 0.001; **** *P* < 0.0001.

To determine whether *rnfC* deletion affects CRC tumor development, we next treated cultured HCT116 cells with various fusobacterial strains at the MOI of 50 for 72 h before imaging and quantification of resulting spheroids. Remarkably, HCT116 cells treated with the parent strain formed larger spheroids than HCT116 cells treated with the *rnfC* mutant, relative to the untreated samples (PBS) (Fig. 6C-6D). Consistent with the role of FadA in tumor development, deletion of *fadA* reduced the ability of fusobacteria to form spheroids (Fig. 6C-6D; Δ*fadA*). To further examine the effect of Rnf on stimulating tumor growth, pre-grown HCT116 spheroids were infected with fusobacteria 48 h before imaging and quantification. Consistent with the above results, the Δ*rnfC* and Δ*fadA* mutants were defective in promoting spheroid development (Fig. 6E-6F). Together, our results demonstrate that the Rnf complex metabolically modulates the expression of FadA and FadA amyloids via its impact on several two-component transduction systems, ultimately promoting fusobacterial pathogenesis as an oncobacterium.

## DISCUSSION

The human oral cavity is home to a rich diversity of microorganisms that must adapt to consistent changes in nutrient and redox availability depending on individual specific factors including age, sex, diet, health and disease status, and microbiome composition. As such, metabolic flexibility and versatility are key to microbial fitness in these communities (38). The oral anaerobe *F. nucleatum* is well-known for its ability to disseminate beyond the oral cavity, colonizing a variety of host tissues of disparate cellular compositions and potentiating several diseases, such as preterm birth (14, 19, 39, 40) and CRC (17, 18, 23, 41). How *F. nucleatum* maintains metabolic plasticity for its virulence capability under these diverse metabolic landscapes is of considerable current interest. To that end, we have recently reported that *F. nucleatum* encodes a functional ferredoxin:NAD^+^ oxidoreductase, or the Rnf complex, which acts as a versatile metabolic exchange center to conserve energy via the production of an ion-motive force (IMF) from metabolism of multiple amino acids (Lys, His, Glu, Gln, Met, Cys). As a result, the Rnf complex expands the redox range of this pathobiont, metabolically stimulating a multitude of pathophysiological traits outside the oral cavity to promote preterm birth in a mouse model of infection (8). Studies reported here reveal that the Rnf complex is also crucial for a second pathogenic trait of *F. nucleatum* – its ability to act as an oncobacterium – and uncover a novel gene regulatory network in which the Rnf complex modulates multiple response regulators to control the cell surface assembly of a key tumor promoting factor – the FadA adhesin (13).

Our studies exploring the function of the Rnf complex in multiple aspects of fusobacterial biology and pathogenesis entailed a systematic physiologic and biochemical comparison of *F. nucleatum* strains that have either an intact *rnf* genetic locus or a targeted in-frame deletion of *rnfC* – the first gene of the *rnf* operon (8). With this approach and comparing both RNA and protein levels in the parent, mutant and complemented strains, we have demonstrated here that the loss of the functional Rnf complex drastically reduces the expression of the FadA adhesin both at the transcriptional and post-translational levels (Fig. 1A-1D). Visualization of FadA on the bacterial surface through immunofluorescence microscopy has revealed that surface FadA is assembled at the mature cell pole (Fig. 1C & 1E), which suggests that FadA surface assembly is independent of nascent peptidoglycan biosynthesis. A key feature of FadA’s physiological function is that it forms amyloid-like fibers on the bacterial surface under high salt conditions (23) that mimics the osmotic environment in the human bloodstream and proximal colon, where the bulk of the electrolytes are absorbed (42, 43). A significant finding we reported here is that the disruption of Rnf greatly diminishes FadA amyloid formation because the FadA amyloids play a significant role in biofilm formation and the proliferation of CRC cells (Fig. 3).

The critical question that arises is how the absence of Rnf function leads to a diminished expression and surface assembly of FadA. In this regard, it is prudent to recall that *F. nucleatum* ATCC 23726 encodes seven TCSs (Table S1), of which some are known to regulate the expression of RadD and Fap2 adhesins and promote extra-oral disease in mouse models of infection (27, 31). Indeed, our qRT-PCR analysis revealed that the disruption of Rnf reduces expression of several response regulators including those coding for CarR, ArlR, and S1, which in turn modulate FadA expression (Fig. 4 & 5). Intriguingly, since the level of *fadA* transcripts remains unchanged in the *carR*, *arlR*, and *s1* mutants (Fig. 4), the reduced surface expression of FadA in these mutants led to the hypothesis that FadA is post-translationally processed by a factor whose expression might be targeted by these response regulators. Because FadA harbors a signal peptide sequence known to be cleaved for FadA secretion and amyloid formation, it is logical to invoke that the signal peptidase LepB is the postulated factor that is involved in the post-translational processing of FadA. Indeed, we show that in the absence of *rnfC*, *arlR*, or *s1*, the level of *lepB* transcript is significantly reduced, and that the targeted reduction of *lepB* transcript, via CRISPR interference, also reduces the surface expression of FadA (Fig. 5). Thus, our results establish that LepB is the major signal peptidase that post-translationally processes FadA and that LepB is transcriptionally controlled by the action of Rnf on the response regulator ArlR and S1.

Given that deletion of *rnfC* reduces the level of *fadA* transcripts (Fig. 1B), which are not significantly affected by deletion of *carR*, *arlR*, or *s1* (Fig. 4C), how does the Rnf complex transcriptionally control *fadA* expression? Since metabolic defects associated with *rnf* mutants negatively impact transcript expression of all 7 response regulators (Fig. 5A), it is possible that *fadA* expression is subject to cross-regulation by multiple response regulators. As such, their individual deletion mutants might not sufficiently affect *fadA* transcripts. Alternatively, metabolic blockage by Rnf deficiency may trigger a transcriptional suppression of *fadA* by a suppressor. Future studies will examine these possibilities.

As FadA is a prominent adhesin of fusobacteria required for fusobacterial colorectal colonization, invasion, and enhancement of CRC (14, 18, 19, 22, 44), it was important to examine how defective FadA expression in Rnf complex-deficient fusobacteria affected these pathogenic processes. We therefore tested the ability of the *rnfC* deletion mutant to adhere to, invade and stimulate tumor formation of human CRC cells, HCT116, *in vitro*. Surprisingly, *rnfC*-deletion had no noticeable impact on CRC cell adhesion (Fig. 6A), which indicates that Fap2-binding to Gal-GalNAc and RadD-binding to CD147 must be sufficient for effective gut colonization by fusobacteria (25, 44). In striking contrast to the normal adherence, CRC cell invasion was severely compromised by the Δ*rnfC* disruption of the Rnf complex (Fig. 6B).

Lastly, we monitored spheroid formation by HCT116 CRC cells in vitro as a model for colorectal tumor formation. Our results clearly demonstrate that Δ*rnfC* mutant fusobacteria are grossly defective in inducing spheroid formation *in vitro* (Fig. 6C – Fig. 6D), nor could the mutant fusobacteria promote growth of pre-formed spheroids (Fig. 6E – Fig. 6F).

In conclusion, the Rnf complex expands the metabolic versatility of *F. nucleatum* through its role in amino acid metabolism, driving several two-component signaling systems to facilitate FadA expression. As such, deletion of *rnfC* impairs this novel gene regulatory network, and thereby greatly hinders fusobacterial invasion of CRC cells and their impact on tumor propagation. Given the high conservation of the Rnf complex in many anaerobic bacterial pathogens, and its absence from eukaryotes, this ancient respiratory enzyme serves as an attractive drug target to combat *F. nucleatum*-associated malignancies.

## MATERIALS AND METHODS

### Bacterial strains, plasmids, and media

All bacterial strains and plasmids used in this study are listed in *Supplemental Material (SM)* Table S2. *F. nucleatum* strains were grown in tryptic soy broth supplemented with 1% Bacto Peptone and 0.25% fresh L-cysteine (TSPC) or on TSPC agar plates at 37°C in an anaerobic chamber as previous described (8). *Escherichia coli* strains were grown in Luria Broth (LB) at 37°C. When required, chloramphenicol, thiamphenicol, or penicillin G were added to the medium at a concentration of 15 µg/mL, 5 µg/mL, or 10 µg/mL, respectively. All reagents were purchased from Sigma-Aldrich unless noted otherwise.

### Plasmid construction

i. pMCSG7-based plasmids: pMCSG7 was used to clone vectors expressing recombinant proteins FadA, CarR, and Fap2 for antibody production, and vectors pMCSG7-FadA, pMCSG7-CarR, and pMCSG7-Fap2 were created with the primer sets using ligation-independent cloning (LIC) as previously reported (45). Briefly, a pair of primers (LIC-FadA-F/R, LIC-CarR-F/R, or LIC-Fap2-F/R listed in *SM* Table S3) was used to amplify part of the coding sequence of *fadA* (corresponding residues 48-129), *carR* (residues 2-224), and *fap2* (residues 3401-3786) from the chromosomal DNA of *F. nucleatum* ATCC 23726. Generated amplicons were inserted into pMCSG7, and the resulting plasmids were introduced into *E. coli* DH5α for propagation and verification by DNA-sequencing. Verified clones were then introduced into *E. coli* BL21 (DE3) for protein expression.
ii. pRnfC: The primer pair com-rnfC-F/R (Table S3) was used to amplify the *rnfC* coding region and its promoter from *F. nucleatum* ATCC 23726 chromosomal DNA. KpnI and NdeI restriction sites were appended to the amplicon, and the PCR product was digested and cloned into pCWU6 as previously described (46). The generated vector was subjected to DNA sequencing for confirmation.
iii. Gene deletion plasmids: pCM-GalK (Table S2) was used to generate vectors for deletion of *fadA*, *arlR*, *cheY*, *eutV*, *s1*, and *ypdB* according to a published protocol (28). Briefly, 1-kb flanking regions upstream and downstream of individual genes of interest were PCR amplified using a specific set of primers (Table S3), and the PCR products were cloned into pCM-GalK. The generated vectors were subjected to DNA sequencing for confirmation.

### Gene deletion in *F. nucleatum*

Using the generated gene deletion plasmids mentioned above (see Table S2), non-polar, in-frame deletion mutants, Δ*fadA*, Δ*arlR*, Δ*cheY*, Δ*ypdB,* Δ*eutV,* and Δ*s1*, were generated as previously described (8, 27, 46).

### Depletion of *F. nucleatum lepB* by CRISPRi

The CRISPRi-based plasmid pZP4C (35) was used to generate pZP4C-lepB (Table S2). For PCR amplification of a single guide RNA (sgRNA) that targets *lepB* (RS05265), primers, Sg-lepB_F and Sg-RNA-R (Table S3), were used with *F. nucleatum* ATCC 23726 chromosomal DNA as template. The generated sgRNA cassette was cloned into pZP4C between MscI and NotI restriction sites. The generated vector was transformed into *E. coli* DH5α for DNA amplification and verification prior to being introduced into ATCC 23726 by electroporation. Transformants were selected on TSPC agar plates containing thiamphenicol (5 µg/ml). Overnight cultures of obtained colonies anaerobically grown in TSPC broth supplemented with thiamphenicol at 37°C were used to inoculate fresh cultures with a starting optical density at 600 nm (OD_600_) of 0.1 in the presence or absence of 2mM theophylline. The resulting cultures were grown in an anaerobic chamber for 18 h prior to being normalized to an OD_600_ of 1.0 for western blotting analysis.

### Western blotting

Expression of fusobacterial proteins were analyzed by immunoblotting with antibodies against FadA (α-FadA; 1:8000), CarR (α-CarR; 1:5000), and Fap2 (α-Fap2; 1:4000). The antibodies were generated using *E. coli* BL21 (DE3) strains harboring pMCSG7-FadA, pMCSG7-CarR, or pMCSG7-Fap2 (mentioned above) as previously described (28, 47). Briefly, cell-free lysates obtained from *E. coli* cell cultures were subjected to protein purification by affinity chromatography. The purified proteins were used for antibody production (Cocalico Biologicals, Inc.). To perform immunoblots, overnight (∼17 h) fusobacterial cultures were harvested and normalized by OD_600_. 1 ml-aliquots of normalized cultures were subjected to protein precipitation by trichloroacetic acid (TCA), followed by acetone wash as previously described (46). Protein samples were suspended in SDS-containing sample buffer with 3M urea, separated by SDS-PAGE using a 4-15% Tris-Glycine gradient gel (Nacalai USA, Inc.), and immunoblotted with specific antibodies. When indicated, band intensity was calculated using ImageJ.

### qRT-PCR

Fusobacterial strains were cultured overnight (∼17 h) and normalized to OD_600_ of ∼2.0. Normalized cells harvested by centrifugation were used to extract total RNA using the RNeasy Mini Kit (Qiagen) according to manufacturer’s instructions and as previously described (8, 28). Approximately 1 µg of purified RNA, free from DNA by treatment with DNase I (Qiagen), was reverse transcribed into cDNA using iScript RT supermix (Bio-Rad) according to the manufacturer’s protocol. Obtained cDNA was used for qRT-PCR with appropriate primers (Table S3) and SYBR Green PCR Master Mix (Bio-Rad). The ΔΔ*C_T_*method was used to calculate fold changes in gene expression between samples. Briefly, ΔΔ*C_T_* = Δ*C_T1_* -Δ*C_T2_*, where Δ*C_T_*= *C_T_* (target) - *C_T_* (housekeeping gene). The *16S* rRNA gene was used as a reference and reactions without reverse transcriptase used as a control to assess genomic DNA contamination.

### Immunofluorescence microcopy

Immunofluorescence microscopy was performed as previously described (8). Circular glass coverslips were placed in a 24-well plate and 0.2 mL aliquots of poly-L-lysine were used to coat the surface for 15 min before washing with sterile water and air-drying for 2 h. Fusobacterial cells grown overnight (∼17 h) with or without 100 mM sodium chloride (NaCl) were harvested by centrifugation and washed twice before being normalized to OD_600_ of ∼ 0.3. Aliquots (∼0.2 mL) of resulting cell suspensions were used to coat the surface of poly-L-lysine-treated glass coverslips and incubated at room temperature for 20 min. Cells were fixed using 2.5% formaldehyde (in PBS) for 20 min, washed with PBS, and blocked for 1 h with 3% wt/vol bovine serum albumin (FadA, Fap2) or 5% skim milk (amyloid), both diluted in PBS supplemented with tween-20 (PBST). Cells were incubated with α-FadA (1:300), α-Fap2 (1:200), or α-Aβ42 (1:500) for 1 h and then AlexaFluor488- (FadA, Fap2) or AlexaFluor594-conjugated (amyloid) goat anti-rabbit IgG (1:200) for another hour in the dark, followed by washing in PBS three times. Coverslips were mounted on glass slides with VECTASHIELD anti-fade mounting medium containing DAPI (Vector Laboratories, Inc.). Images were taken using a fluorescence microscope (Keyence BZ-X800) and fluorescent units normalized and quantified using ImageJ.

For cell labeling with fluorescently-tagged D-amino acids (FDAAs), the experiment was performed as previously described (29, 48), with some modification. Briefly, overnight cultures of fusobacterial strains were used to inoculate fresh cultures normalized to OD_600_ of 0.05 in TSPC containing 0.5 mM HADA (Bio-Techne). Cells were grown for 14 h, harvested by centrifugation, washed in PBS, and inoculated in fresh cultures normalized to OD_600_ of 0.3 in TSPC containing 0.5 mM Rf470DL (Bio-Techne). Cells were grown for 3 h before being washed twice in PBS. The resulting cell suspensions were used for labeling with α-FadA and AlexaFluor488 as described above.

### Adherence and invasion of colorectal cancer cells

Adherence and invasion assays were performed as previously described (31), with minor modifications. Human colorectal cancer cells, HCT116 (American Type Culture Collection), were grown in Dulbecco’s modification of Eagle’s medium (DMEM) supplemented with 10% FBS and 1% penicillin G in 24-well tissue culture-treated plates. HCT116 cells cultured to 80% confluency were washed with DMEM supplemented with 10% FBS to remove penicillin G and infected at an MOI (multiplicity of infection) of 50 with indicated fusobacterial strains grown to mid-exponential phase in TSPC. For adherence, fusobacteria were allowed to adhere to HCT116 CRC cells for 4 h before gently washing twice with PBS to remove unattached fusobacterial cells. For invasion, a similar procedure was employed, except that after 3 h of infection, HCT116 CRC cells were treated with PBS supplemented with gentamicin (200 µg/mL) for 1 h to kill all extracellular fusobacteria, followed by washing twice in PBS. To enumerate fusobacterial cells, HCT116 cells were lysed in distilled water for 10 min, followed by serial dilution on TSPC plates for CFU counts.

### Formation and growth of CRC spheroid tumors

HCT116 cells were grown in DMEM supplemented with 10% FBS and 1% penicillin G to 80% confluency and seeded into 24-well ultra-low attachment plates. For spheroid formation, HCT116 cells were immediately infected with fusobacterial strains grown to mid-exponential phase at an MOI of 50 for 72 h. For measurement of spheroid growth, mammalian cells were allowed to grow for 24 h before challenging with fusobacterial strains grown to mid-exponential phase at an MOI of 50 for 48 h. Resulting spheroids were imaged using phase-contrast microscopy and their surface area quantified using ImageJ.

## ACKNOWLEDGMENTS

We thank our lab members for their discussion and critical review of the manuscript. Research reported in this publication was supported by the National Institute of Dental & Craniofacial Research (NIDCR) of the National Institutes of Health (NIH) under Award Numbers DE026758 and DE033900 (to H.T.-T). T.A.B. was supported by the UCLA Dentist-Scientist and Oral Health-Researcher Training Program, NIDCR Grant T90DE030860, and the UCLA Eugene V. Cota Robles fellowship. The content is solely the responsibility of the authors and does not necessarily represent the official views of the NIH.

## AUTHOR CONTRIBUTIONS

T.A.B. and H.T.-T. designed research; T.A.B., J.H.L., C.C., Wu C, B.A.H., and R.M.A., performed research; T.A.B., A.D., and H.T.-T. analyzed data; and T.A.B., A.D., and H.T.-T. wrote the paper.

